# Hematopoietic Stem Cells Undergo Differentiation State Reprogramming to Overcome Venetoclax Sensitivity in Patients with Myelodysplastic Syndromes

**DOI:** 10.1101/2023.09.25.559439

**Authors:** Juan Jose Rodriguez-Sevilla, Irene Ganan-Gomez, Feiyang Ma, Kelly Chien, Guillermo Montalban-Bravo, Rashmi Kanagal-Shamanna, Sanam Loghavi, Vera Adema, Alexandre Bazinet, Guillermo Garcia-Manero, Simona Colla

## Abstract

While the molecular mechanisms of acute myeloid leukemia failure to venetoclax-based therapy have been recently clarified, the mechanisms whereby patients with myelodysplastic syndromes (MDS) acquire secondary resistance to venetoclax after an initial response remain to be elucidated.

Here, we show for the first time that MDS hematopoietic stem cells (HSCs) can undergo hierarchical differentiation reprogramming toward a granulo-monocytic-biased transcriptional state through the acquisition or expansion of clones with an isolated trisomy 8 cytogenetic aberration and *STAG2* or *RUNX1* mutations. This hierarchical rewiring changes HSCs’ survival dependence from BCL-2-mediated anti-apoptotic pathways to TNFα-induced pro-survival NF-κB signaling and overcomes venetoclax-mediated cytotoxic effect. These findings underscore the importance of close molecular monitoring of patients with MDS enrolled in clinical trials of venetoclax to prevent HSC transcriptional reprogramming before the disease becomes resistant to this therapy.

## MAIN

The hematopoietic stem cell (HSC) hierarchy of myelodysplastic syndromes (MDS) predicts the biological mechanisms of progression after failure to frontline hypomethylating agents (HMAs) and can guide the design or choice of second line therapeutic approaches^1^. We previous showed that, compared with those with a “granulocytic-monocytic progenitor (GMP) pattern” of differentiation, MDS patients with an immunophenotypic “common myeloid progenitor (CMP) pattern” of differentiation who received venetoclax-based therapy had a shorter cumulative time to complete remission and a longer recurrence-free survival duration, primarily because venetoclax can efficiently target only “CMP pattern” HSCs, whose survival depends on BCL2-^1^.

Although confirmed in a larger cohort of samples (n = 28; 12 “CMP pattern” MDS and 16 “GMP pattern” MDS) (Supplemental Figures 1a,b and Supplemental Table 1), our longer-term follow-up survival of MDS patients (median: 18.2 months) enrolled in venetoclax-based clinical trials showed that those with “CMP pattern” MDS eventually lose response and/or progress to acute myeloid leukemia (AML) after an initial disease remission (n=6 of 8 “CMP pattern” patients with an initial response), which suggests that alternative approaches are needed for these patients who would otherwise have no other therapeutic options.

Here, to dissect the cellular and molecular mechanisms of secondary venetoclax-based therapy failure, we performed multi-omics analyses of sequential samples from 4 “CMP pattern” MDS patients whose initial disease response to venetoclax-based therapy was associated with HSC depletion (Supplemental Figure 1c and Supplemental Table 2).

These analyses showed that the “CMP pattern” immunophenotypic architecture (Supplemental Figure 1d) and the hematopoietic stem and progenitor cell (HSPC) transcriptomic signature (Supplemental Figures 1e,f) persisted at disease recurrence in patients with *TP53* mutations (2 out of 4), which is consistent with previous findings that *TP53* mutations confer an intrinsic resistance to BCL-2 inhibition^2^.

However, the HSPC hierarchy switched to “GMP pattern” MDS in the other 2 patients (UPN#1 and UPN#2) before venetoclax failure (Figures 1a,b and Supplemental Figures 1g,h). This immunophenotypic hierarchical change was associated with the acquisition or expansion of clones with an isolated trisomy 8 cytogenetic aberration and “GMP pattern”–prone^1^ *STAG2*- or *RUNX1*-mutations (Figure 1c and Supplemental Figure 1i,j). Single cell RNA-sequencing (scRNA-seq) analyses of mononuclear cells (MNCs) from sequential bone marrow (BM) samples from these 2 patients (Figure 1d and Supplemental Figure 1k) confirmed that HSCs were significantly depleted during disease remission but expanded at therapy failure (Supplemental Figure 1l). Differential expression analyses of sequential BM samples collected during different disease stages showed that the acquisition of *STAG2*- or *RUNX1*-mutant clones with trisomy 8 not only rewired MDS HSCs’ differentiation state towards a myeloid-biased transcriptional signature (Supplemental Figure 1m) but also changed HSCs’ survival dependence from BCL-2-mediated anti-apoptotic pathways to TNFα-induced pro-survival NF-κB signaling, thus enabling HSCs to evade the cytotoxic effects of venetoclax (Figure 1e and Supplemental Figures 1n-p).

**Figure 1.**
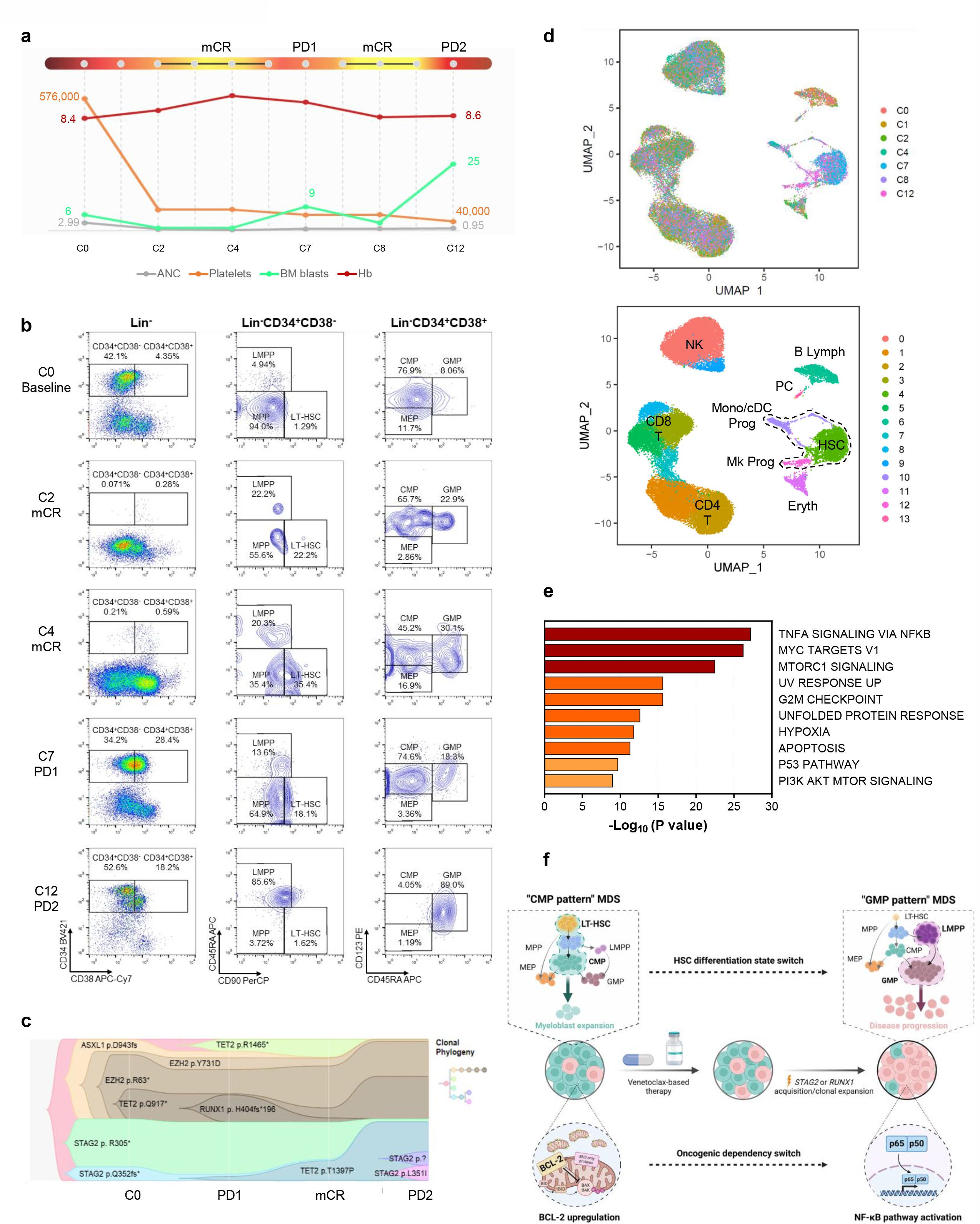
Mutation-induced MDS HSCs’ transcriptional reprogramming overcomes venetoclax-based therapy vulnerability. **(a)**Schematic of UPN#1’s clinical course. After HMA therapy failure (cycle 0 [C0]), UPN#1 received 5-azacitidine (75 mg/m^2^ for 5 days) and venetoclax (400 mg/m^2^ for 14 days) every month. The patient had mCR at cycle 2 (C2), however, after the venetoclax dose was reduced to 100 mg/m^2^, the patient had an initial disease progression (PD1) at cycle 7 (C7). The patient had mCR after the venetoclax dose was increased to 200 mg/m^2^ at cycle 8 (C8) but had progression to AML (PD2) at cycle 12 (C12). Hb, hemoglobin; ANC, absolute neutrophil count. Units: blasts, %; Hb, g/dL; ANC, ×10^9^/L; platelets, ×10^9^/L. **(b)**Flow cytometry plots of lineage (Lin)^-^CD34^+^CD38^-^ HSCs and Lin^-^CD34^+^CD38^+^ myeloid hematopoietic progenitor cells in the BM of UPN#1 at sequential timepoints before and during venetoclax-based therapy. LT-HSC, long-term hematopoietic stem cells; MPP, multipotent progenitors; LMPP, lymphoid-primed multipotent progenitors; CMP, common myeloid progenitors; GMP, granulocytic-monocytic progenitors; MEP, megakaryocyte erythroid progenitors. **(c)**Fish plot of the clonal evolution pattern inferred from NGS data. The phylogenetic trees show the estimated order of mutation acquisition and the proportion of subclones with different combinations of mutations at each timepoint. In UPN#1, clonal evolution was associated with the immunophenotypic HSPC hierarchical change and the acquisition of 2 *STAG2* mutations. **(d)**UMAP plots of scRNA-seq data from BM MNCs isolated from patient UPN#1 (n=39,206). Each dot represents 1 cell. Different colors indicate the sample origin (top) and cluster identity (bottom). HSC, hematopoietic stem cell; Mk, megakaryocytic; Mono, monocytic; cDC, classic dendritic; Prog, progenitor; Eryth, erythroblast; NK, natural killer; Lymph, lymphocyte; PC, plasma cell. Dotted lines indicate the HSPC compartment. **(e)** Pathway enrichment analysis of the genes that were significantly upregulated in HSCs from UPN#1 (cluster 4 in Figure 1d) at the time of PD2 compared with those in HSCs at the time of PD1 (*P adj*≤0.05). The top 10 Hallmark gene sets are shown. **(f)** Proposed working model. After an initial response to venetoclax-based therapy, the acquisition or expansion of trisomy 8–harboring clones with *STAG2* or *RUNX1* mutations reprograms the HSPC hierarchy and switches these cells’ dependence from BCL-2-to NF-κB– mediated survival programs, which leads to secondary failure to venetoclax-based therapy.

Notably, trisomy 8 was also significantly associated with *STAG2* mutations (*P*=0.03) and conferred shorter duration of response to venetoclax-based therapy regardless of prior treatment in patients with “CMP pattern” MDS but not those with “GMP pattern” MDS (n=53 patients treated with venetoclax-based therapies for whom immunophenotypic data were available) (Supplemental Figure 1q ad Supplemental Table 3), which suggests that trisomy 8 is a predictive biomarker of venetoclax resistance in patients with “CMP pattern” MDS.

Together, these results reveal a novel mechanism of venetoclax-based therapy failure. “CMP pattern” HSCs under venetoclax therapy undergo survival pressure, which results in the acquisition or expansion of clones carrying specific genetic alterations that change these cells’ dependence on BCL2-mediated antiapoptotic pathways for survival (Figure 1e).

Our study suggests that MDS patients receiving venetoclax-based therapy should be monitored closely for the acquisition or expansion of trisomy 8–harboring clones with *STAG2* or *RUNX1* mutations and enrolled in clinical trials of agents targeting NF-κB signaling effectors before their disease undergoes HSC transcriptional reprogramming and becomes resistant to venetoclax.

## METHODS

### Human primary samples and clinical data analysis

We analyzed MDS patients who received venetoclax-based therapy at MD Anderson Cancer Center. Patients were enrolled in 1 of 3 phase I/II clinical trials (NCT04160052^3^, NCT04550442^4^, or NCT04655755^5^). Patient characteristics, laboratory values, and BM data, including cytogenetics and next-generation sequencing (NGS) data, were assessed at baseline (before venetoclax-based therapy) and thereafter as clinically warranted. Genomic DNA was extracted from whole BM aspirates and subjected to 81-gene target polymerase chain reaction–based sequencing using an NGS platform as described previously^6^. Testing was performed in a Clinical Laboratory Improvement Amendments– certified laboratory. Risk stratification was performed using the Revised International Prognostic Scoring System (IPSS-R), and MDS was classified as lower-risk (IPSS-R score ≤3.5) or higher-risk (IPSS-R score >4) MDS^7,8^. Disease response was categorized according to the International Working Group 2006 criteria for MDS, and patients with responsive disease included those with complete response (CR), marrow CR (mCR), hematologic improvement (HI), or a combination of mCR and HI^9^. Response duration was defined as the time from first documented response to first documented disease progression or death, whichever occurred first. To evaluate the mechanisms of secondary failure to venetoclax-based therapy, we analyzed 28 MDS patients enrolled in the 3 clinical trials in whom HMA therapy had failed. To evaluate the impact of trisomy 8 on the survival of MDS patients treated with venetoclax-based therapy, we analyzed the clinical data of 53 patients who were enrolled in the 3 clinical trials regardless of prior therapies and for whom immunophenotypic data were available.

Samples were obtained in accordance with the Declaration of Helsinki from MD Anderson’s Department of Leukemia under protocol PA15-0926 with the approval of the corresponding Institutional Review Boards. Written informed consent was obtained from all donors, and all diagnoses were confirmed by dedicated hematopathologists. The clinical characteristics of the patients included in this study are shown in Supplemental Tables 1-4. MNCs were isolated from each sample using the standard gradient separation approach with Ficoll-Paque PLUS (GE Healthcare Lifesciences, Pittsburgh, PA).

### Flow cytometry and fluorescence-activated cell sorting (FACS)

Quantitative flow cytometric analyses and FACS of human live MNCs were performed using a previously described gating strategy and antigen panel^1,10^ and antibodies against CD2 (RPA-2.10; 1:20), CD3 (SK7; 1:10), CD14 (MφP9; 1:20), CD19 (SJ25C1; 1:10), CD20 (2H7; 1:10), CD34 (581; 1:20), CD56 (B159; 1:40), CD123 (9F5; 1:20), and CD235a (HIR2; 1:40; all from BD Biosciences, Franklin Lakes, NJ); CD4 (S3.5; 1:20), CD11b (ICRF44; 1:20), CD33 (P67.6; 1:20), and CD90 (5E10; 1:10; all from Thermo Fisher Scientific, Waltham, MA); CD7 (6B7; 1:20) and CD38 (HIT2; 1:20; both from BioLegend, San Diego, CA); CD10 (SJ5-1B4; 1:20; Leinco Technologies, St Louis, MO); and CD45RA (HI100; 1:10; Tonbo Biosciences, San Diego, CA).

FACS-purified samples were acquired with a BD Influx Cell Sorter (BD Biosciences), and the cell populations were analyzed using FlowJo software (version 10.7.1, Ashland, OR). All experiments included single-stained controls and were performed at MD Anderson’s South Campus Flow Cytometry and Cellular Imaging Facility.

### scRNA-seq

scRNA-seq was performed as we described previously^1^. Briefly, FACS-purified live BM MNCs were prepared and sequenced at MD Anderson’s Advanced Technology Genomics Core. Sample concentration and cell suspension viability were evaluated using a Countess II FL Automated Cell Counter (Thermo Fisher Scientific) and manual counting.

Samples were normalized for input onto the Chromium Single Cell A Chip Kit (10x Genomics, Pleasanton, CA), in which single cells were lysed and barcoded for reverse-transcription. The pooled single-stranded, barcoded cDNA was amplified and fragmented for library preparation. Pooled libraries were sequenced on a NovaSeq6000 SP 100-cycle flow cell (Illumina, San Diego, CA).

The sequencing analysis was carried out using 10X Genomics’ CellRanger software (version 3.0.2). Fastq files were generated using the CellRanger MkFastq pipeline (version 3.0.2). Raw reads were mapped to the human reference genome (refdata-cellranger-GRCh38-3.0.0) using the CellRanger Count pipeline. Multiple samples were aggregated using the Cellranger Aggr pipeline. The digital expression matrix was analyzed with the R package Seurat (version 3.0.2)^11^ to identify different cell types and signature genes for each. Cells with fewer than 500 unique molecular identifiers or greater than 50% mitochondrial expression were removed from further analysis. The Seurat function NormalizeData was used to normalize the raw counts. Variable genes were identified using the FindVariableFeatures function. The ScaleData function was used to scale and center expression values in the dataset, and the number of unique molecular identifiers was regressed against each gene. Uniform manifold approximation and projection (UMAP) was used to reduce the dimensions of the data, and the first 2 dimensions were used in the plots. The FindClusters function was used to cluster the cells. Marker genes for each cluster were identified using the FindAllMarkers function. Cell types were annotated based on the marker genes and their match to canonical markers^12,13^. Pathway analyses of differentially expressed genes were conducted using Metascape^14^. The GMP enrichment score was calculated based on a previously validated GMP expression signature^15^.

Statistical analyses were performed using R (version 4.0.320), Jamovi (version 2.0.021), and GraphPad (version 9.0.0, San Diego, CA). Two-tailed Student t-test or Mann−Whitney test, as appropriate, and chi-square test were used to compare continuous and categorical variables, respectively. Mutations with variant allele frequency values below 2% were excluded from the plot to model clonal evolution. A comprehensive summary of the mutations for UPN#1 and UPN#2 at every timepoint is provided in Supplemental Table 4. Fish plot visualization was performed using the timescape package (version 3.14) in R (version 4.2.2). The graphical abstract was made using BioRender.

## Data availability

The datasets generated using scRNA-seq are accessible at GEO under GSE241417.

## ACKNOWLEDGMENTS

This work was supported by philanthropic contributions to MD Anderson’s AML and MDS Moon Shot Program, by the National Institutes of Health (NIH) through MD Anderson’s Leukemia SPORE grant (P50 CA100632), and by the Edward P. Evans Foundation. S.C. is a Scholar of the Leukemia and Lymphoma Society. This work used MD Anderson’s South Campus Flow Cytometry and Cellular Imaging Facility and Advanced Technology Genomics Core, both of which are supported in part by the NIH through MD Anderson’s Cancer Center Support Grant (P30 CA16672).

## AUTHOR CONTRIBUTIONS

S.C. designed the research; J.J.R.-S. and I.G.-G. guided the research; J.J.R.-S., I.G.-G., and V.A. performed experiments; J.J.R.-S., K.C., G.M.-B., and A.B. analyzed the clinical data; R.K.-S. and S.L. analyzed the genomic data; F.M. analyzed the scRNA-seq data; J.J.R.-S. and I.G.-G. analyzed the flow cytometry data; G.G.-M. made critical intellectual contributions throughout the project; S.C. wrote the manuscript.

## COMPETING INTERESTS

G.G.-M. reports clinical funding from AbbVie and Amgen. All other authors report no competing interests relative to this work.

Correspondence and requests for materials should be addressed to.

## FIGURE LEGENDS

**Supplemental Figure 1.**
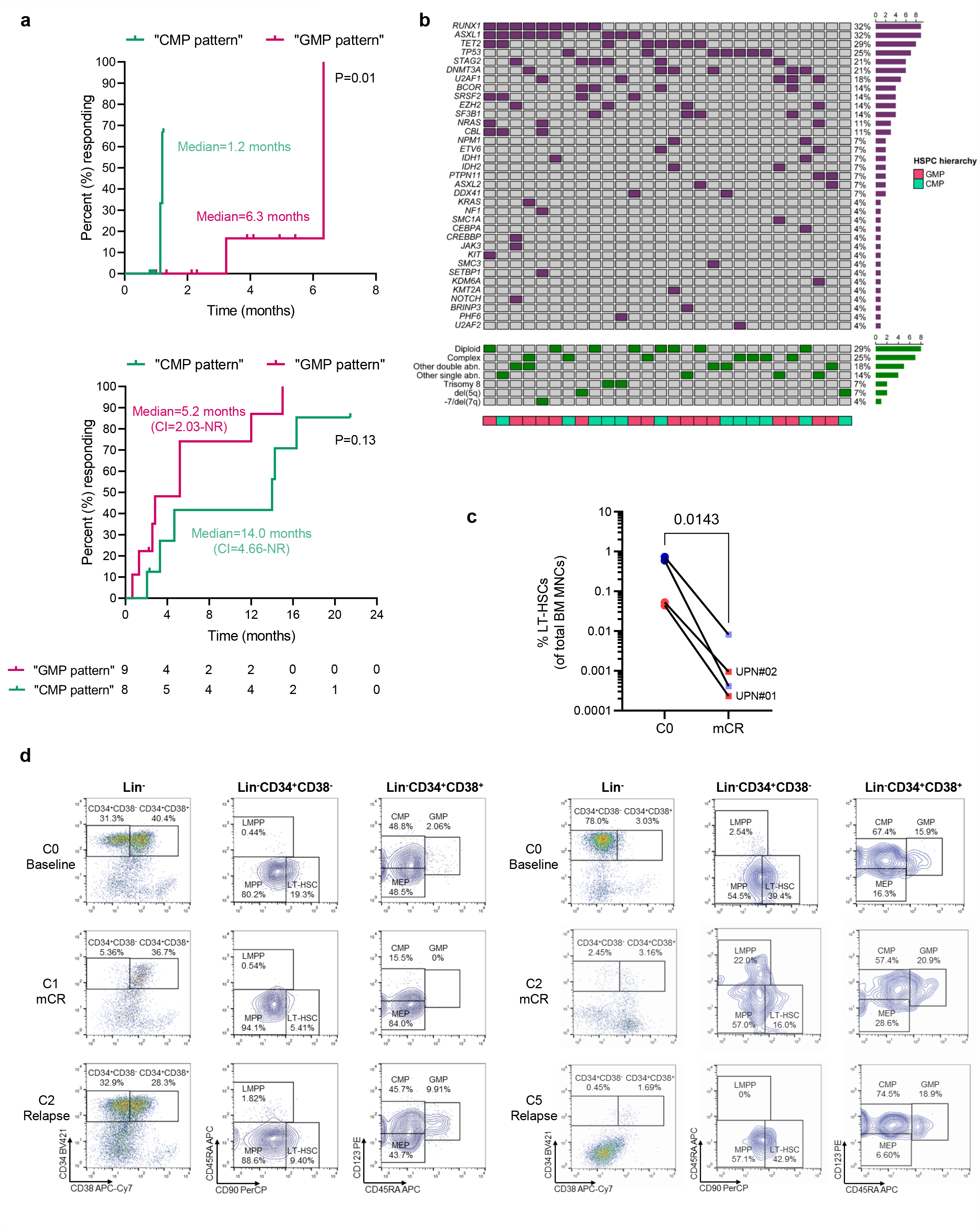
Mutation-induced MDS HSCs’ transcriptional reprogramming overcomes venetoclax-based therapy vulnerability. **(a)**Cumulative times to complete remission (top) and cumulative incidences of relapse for “CMP pattern” (n=8) and “GMP pattern” (n=9) MDS patients whose disease progressed after HMA therapy failure and who were enrolled in clinical trials of venetoclax-based therapy. Statistical significance was calculated using the log-rank (Mantel–Cox) test. **(b)**Oncoplot of molecular alterations (top), cytogenetic alterations (middle), and hierarchical HSPC patterns (bottom) in 28 MDS patients who were enrolled in clinical trials of venetoclax-based therapy after HMA therapy failure. **(c)**Frequencies of LT-HSCs in BM MNCs at the time of HMA therapy failure (cycle 0 [C0]) and at the time of mCR in sequential samples from 4 “CMP pattern” MDS patients for whom HSPC immunophenotypic data were available and whose disease initially responded to venetoclax-**(d)** based therapy but then failed therapy or progressed to AML. Statistically significant differences were determined using a two-tailed Student t-test. **(e)** Flow cytometry plots of lineage (Lin)^-^CD34^+^CD38^-^ HSCs and Lin^-^CD34^+^CD38^+^ myeloid hematopoietic progenitor cells from the 2 “CMP pattern” patients with *TP53* mutations at sequential timepoints before and during venetoclax-based therapy. MPP, multipotent progenitors; LMPP, lymphoid-primed multipotent progenitors; CMP, common myeloid progenitors; GMP, granulocytic-monocytic progenitors; MEP, megakaryocyte erythroid progenitors. **(e)** UMAP plots of the scRNA-seq data for BM MNCs isolated from 1 of the 2 patients with *TP53* mutations (n=10,774; Supplemental Table 2). Each dot represents 1 cell. Different colors indicate the sample origin (top) and cluster identity (bottom). HSC, hematopoietic stem cell; Eryth, erythroblast. Dotted lines indicate the HSPC compartment. **(f)** Pathway enrichment analysis of the genes that were significantly upregulated in HSPCs from 1 of the patients with *TP53* mutations (cluster 8 in Supplemental Figure 1e) at the time of relapse compared with those in HSPCs at the time of HMA therapy failure (C0; *P adj*≤0.05). Hallmark gene sets are shown. **(g)** Schematic of UPN#2’s clinical course. After HMA therapy failure (C0), UPN#2 received 5-azacitidine (75 mg/m^2^ for 5 days) and venetoclax (70 mg/m^2^ for 7 days) every month. The patient had mCR at cycle 2 (C2) but lost response at cycle 9 (C9). Hb, hemoglobin; ANC, absolute neutrophil count. Units: blasts, %; Hb, g/dL; ANC, ×10^9^/L; platelets, ×10^9^/L. **(h)** Flow cytometry plots of Lin^-^CD34^+^CD38^-^ HSCs and Lin^-^CD34^+^CD38^+^ myeloid hematopoietic progenitor cells in the BM of UPN#2 at sequential timepoints before and during venetoclax-based therapy. MPP, multipotent progenitors; LMPP, lymphoid-primed multipotent progenitors; MEP, megakaryocyte erythroid progenitors. **(i)** Fish plot of clonal evolution pattern inferred from NGS data. The phylogenetic trees show the estimated order of mutation acquisition and the proportion of subclones with different combinations of mutations at each timepoint. In UPN#2, clonal evolution was associated with the immunophenotypic HSPC hierarchical change and the acquisition of 2 *RUNX1* mutations. **(j)** Cytogenetic analyses of isolated trisomy 8 in sequential BM samples from patients UPN#1 (top) and UPN#2 (bottom). The frequencies of metaphases with trisomy 8 at C0, mCR, and progression to AML (PD2) or relapse, respectively, are shown. **(k)** UMAP plots of scRNA-seq data for BM MNCs isolated from UPN#2 (n=12,703). Each dot represents 1 cell. Different colors indicate the sample origin (top) and cluster identity (bottom). HSC, hematopoietic stem cell; MPP, multipotent progenitor; LMPP, lymphoid-primed multipotent progenitor; Mk, megakaryocytic; Prog, progenitor; Prolif, proliferative; Mono, monocyte; NK, natural killer; Eryth, erythroblast; Lymph, lymphocyte. **(l)** Cluster distribution of cells in the HSPC compartment from UPN#1 (left) and UPN#2 at the time of HMA therapy failure (C0) and at the time of mCR and PD2 or relapse, respectively, after venetoclax-based therapy. **(m)** Distribution of the enrichment scores for the GMP gene signature in HSCs from UPN#1 (cluster 4 in Figure 1d; top) and those from UPN#2 (cluster 12 in Supplemental Figure 1h; bottom) during the indicated cycles of venetoclax-based therapy. **(n)** Dot plot of the expression levels of genes involved in TNFα signaling through the NF-κB pathway that were significantly (*P adj*≤0.05) upregulated in HSCs (cluster 4) from UPN#1 at cycle 12 (C12; PD2) compared with those in HSCs at cycle 7 (C7; PD1). **(o)** Pathway enrichment analysis of the genes that were significantly upregulated in HSCs from UPN#2 (cluster 12 in Supplemental Figure 1k) at the time of relapse (C9) compared with those in HSCs at the time of HMA therapy failure (C0; *P adj*≤0.05). The top 10 Hallmark gene sets are shown. **(p)** Dot plot of the expression levels of genes involved in TNFα signaling through the NF-κB pathway that were significantly (*P adj*≤0.05) upregulated in HSCs (cluster 12) from UPN#2 at cycle 9 (C9; relapse) compared with those at HMA therapy failure (C0). **(q)** Kaplan-Meier plot of the response durations of “CMP pattern” (top) and “GMP pattern” (bottom) MDS patients enrolled in clinical trials of venetoclax-based therapy based on the presence of trisomy 8. Statistical significance was calculated using the log-rank (Mantel–Cox) test.

---

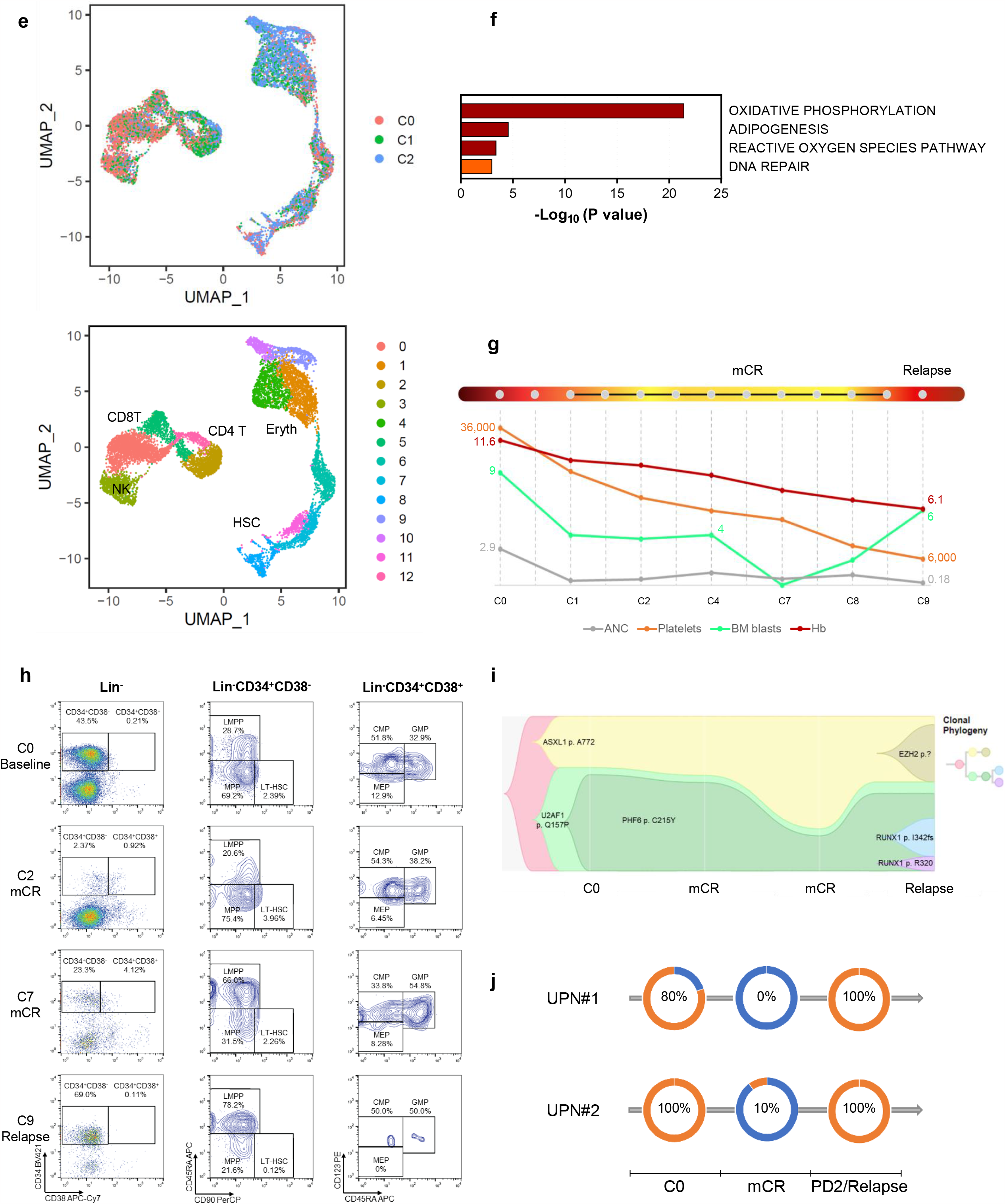

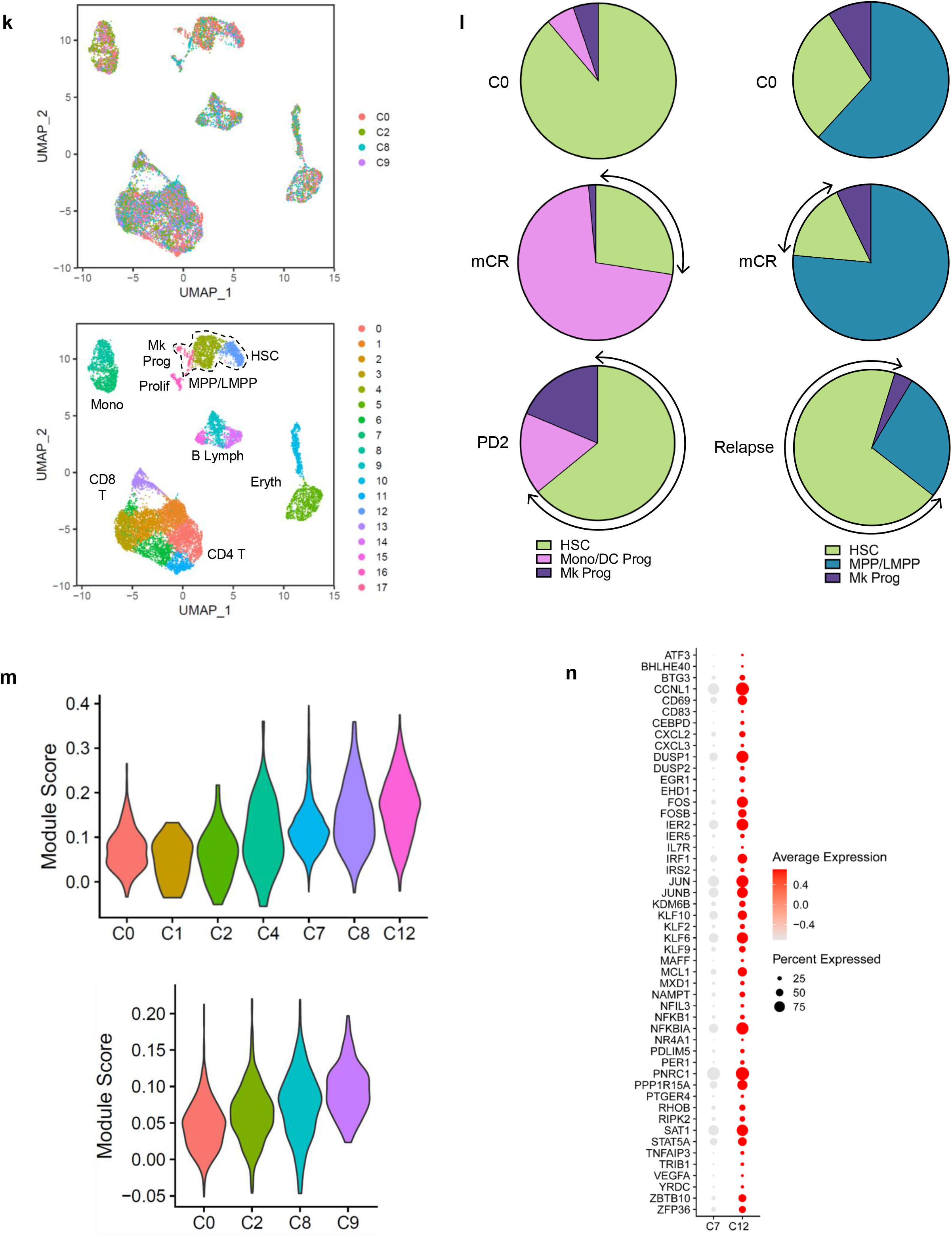

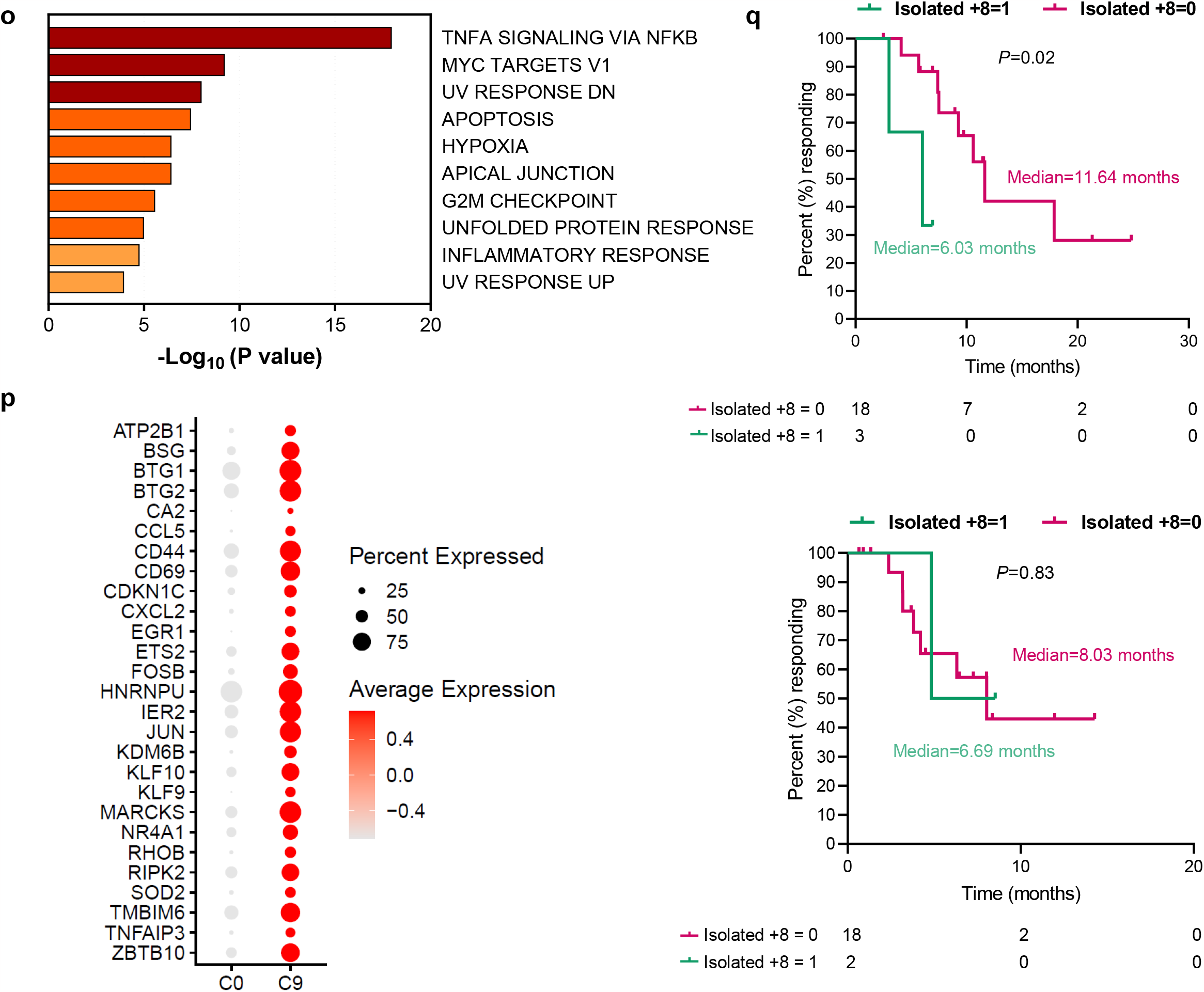

